# Multimodal spatial-omics reveal the heterogeneity and intercellular network characteristics of papillary craniopharyngiomas

**DOI:** 10.64898/2026.08.20.746031

**Authors:** Yu Jiang, Hao Luo, Hui Zheng, Chengxi Li, Xiaofei Wang, Chenyang Tao, Chao Li, Xin Zan, Jianguo Xu, Yaohui Chen

## Abstract

Despite significant advancements in microsurgical techniques in recent years, the treatment and prognosis of craniopharyngiomas remain unsatisfactory. As a central nervous system tumor located adjacent to important brain structures such as the hypothalamus-pituitary axis and accompanied by a highly inflammatory microenvironment, the tumor heterogeneity and tumor microenvironment characteristics of papillary craniopharyngiomas (PCPs) remain unclear. In this study, we integrated multimodal single-cell and spatial profiling from PCP tissue and peripheral blood mononuclear cells (PBMCs) to elucidate the tumor heterogeneity and microenvironment characteristics of PCP. Our single-cell and spatial analyses defined four specific tumor cell states in PCP, representing specific transcriptional regulatory programs and spatial heterogeneity characteristics during tumor progression. By constructing a spatial niche composed of tumor, immune, and stromal cells, we analyzed the cellular and spatial ecosystem of PCP at multiple levels to further assess the communication relationships between different tumor cell states and microenvironment cells. This study established a multidimensional molecular atlas of PCP from the perspectives of cell state, spatial structure, and microenvironment interactions, providing a foundation for understanding its biological behavior and exploring new intervention strategies.

## Introduction

Craniopharyngiomas (CPs) are rare, histologically low-grade epithelial tumors that arise along the craniopharyngeal duct in the sellar and suprasellar regions. Their proximity to the hypothalamus, pituitary stalk, optic apparatus, and floor of the third ventricle creates disproportionate clinical morbidity, including neuroendocrine dysfunction, visual impairment, cognitive deficits, hypothalamic obesity, and sleep disturbance^1,2^. The 2021 World Health Organization (WHO) classification recognizes two biologically distinct entities, adamantinomatous craniopharyngioma (ACP) and papillary craniopharyngioma (PCP)^3^. ACP predominates in children and is characterized by palisading epithelium, stellate reticulum, wet keratin, calcification, and cystic change^4^. Somatic CTNNB1 mutations and aberrant WNT/β-catenin signaling define this subtype and generate tumor cell clusters with nuclear β-catenin accumulation^5^. By contrast, PCP occurs almost exclusively in adults and typically comprises mature non-keratinizing squamous epithelium with papillary architecture, without the wet keratin and prominent calcification of ACP. More than 90% of PCPs harbor the BRAF V600E driver mutation, which sustains MAPK pathway activation^6,7^.

This concentrated molecular dependency creates an unusual opportunity for precision therapy. Historically, PCP has been managed with surgical resection followed, when indicated, by radiotherapy^8^. Yet the close relationship between tumor and the hypothalamic-pituitary axis or optic pathways makes aggressive resection a trade-off between tumor control and irreversible neurological or endocrine injury, whereas conservative resection can leave residual disease at risk of progression or recurrence^9^. The discovery of BRAF V600E in PCP transformed this therapeutic landscape. Combined BRAF and MEK inhibition has produced rapid and profound tumor regression, first in heavily pretreated disease and subsequently in a phase 2 study of newly diagnosed tumors^10^. These responses establish PCP as a compelling model of genomically guided therapy in the central nervous system. However, its broader cellular biology and microenvironmental dependencies remain less well resolved than its dominant oncogenic driver.

A relatively uniform driver genotype does not imply a uniform tumor ecosystem. PCP cells may occupy distinct proliferative, differentiated, stem-like, or stress-associated states, each embedded within immune and stromal compartments. Recent single-cell studies have begun to reveal divergent epithelial programs, macrophage states, and intercellular signaling networks across craniopharyngioma subtypes^11,12^. More broadly, single-cell analyses of human tumors have established that non-genetic malignant cell states can coexist and evolve within a shared genomic background^13^. Nevertheless, the spatial organization of PCP cell states, the cellular neighborhoods that support them, and their interactions with immune and stromal populations remain incompletely defined. Resolving these relationships is necessary to move beyond a mutation-centered view of PCP toward a tissue-level model of the disease.

Bulk molecular profiling averages signals across mixed cell populations and therefore cannot assign molecular programs to their cellular sources or recover the spatial context in which those programs operate^14^. Single-cell RNA sequencing (scRNA-seq) overcomes the first limitation by resolving transcriptional heterogeneity at cellular resolution^13,15,16^, but tissue dissociation removes cells from their native neighborhoods. Spatial transcriptomic methods address this gap by measuring gene expression in intact tissue sections, while imaging-based approaches extend spatial profiling toward single-cell and subcellular resolution^17^.

Complementary multiplexed protein imaging technologies such as co-detection by indexing (CODEX) quantify dozens of markers in situ and reconstruct cell identities, tissue architecture, and cellular neighborhoods^18^. Integrating these modalities can therefore connect molecular cell states to spatial niches and cell-cell interactions that are inaccessible to either bulk or dissociation-based assays alone.

Here, we establish a multimodal PCP cohort that integrates scRNA-seq, Xenium single-cell spatial transcriptomics, CODEX multiplexed spatial proteomics, and complementary bulk omics. We first define the tumor, lymphoid, myeloid, and stromal compartments and resolve their constituent cell states by scRNA-seq. We then map these populations in serial tissue sections using Xenium and CODEX, enabling cross-platform assessment of spatial cellular composition and neighborhood organization. By linking cell state, tissue architecture, and microenvironmental interactions, this study provides a multidimensional molecular atlas of PCP and a framework for identifying cellular dependencies that may complement BRAF-directed therapy.

## Result

To comprehensively characterize the cellular composition, spatial organization, and molecular heterogeneity of papillary craniopharyngioma (PCP), we assembled a multi-omics cohort comprising 77 PCP patients. Multi-omics molecular profiling was performed at the single-cell, spatial, and bulk levels, encompassing transcriptomics, whole-exome sequencing (WES), four-dimensional data independent acquisition (4D-DIA) proteomics, and metabolomics. In this study, we performed single-cell RNA sequencing (scRNA-seq) on tumor tissues from 14 PCP patients, together with three matched PBMC samples. Following stringent quality control and filtering, 146,184 single-cell transcriptomes were retained and integrated to construct a comprehensive cellular atlas of PCP (Fig. 1A). Unsupervised clustering of the scRNA-seq data initially identified four major cellular compartments, comprising tumor cells, lymphoid cells, myeloid cells, and stromal cells (Fig. 1A). We subsequently refined the clustering and annotation of each major compartment, identifying 34 immune cell subsets and seven stromal cell subsets. These included several cell populations with canonical molecular features, such as FLT3^+^ conventional dendritic cells (cDCs), GZMK^+^ CD8^+^ T cells, and regulatory T cells (Tregs) (Fig. 1B–C).

**Figure 1.**
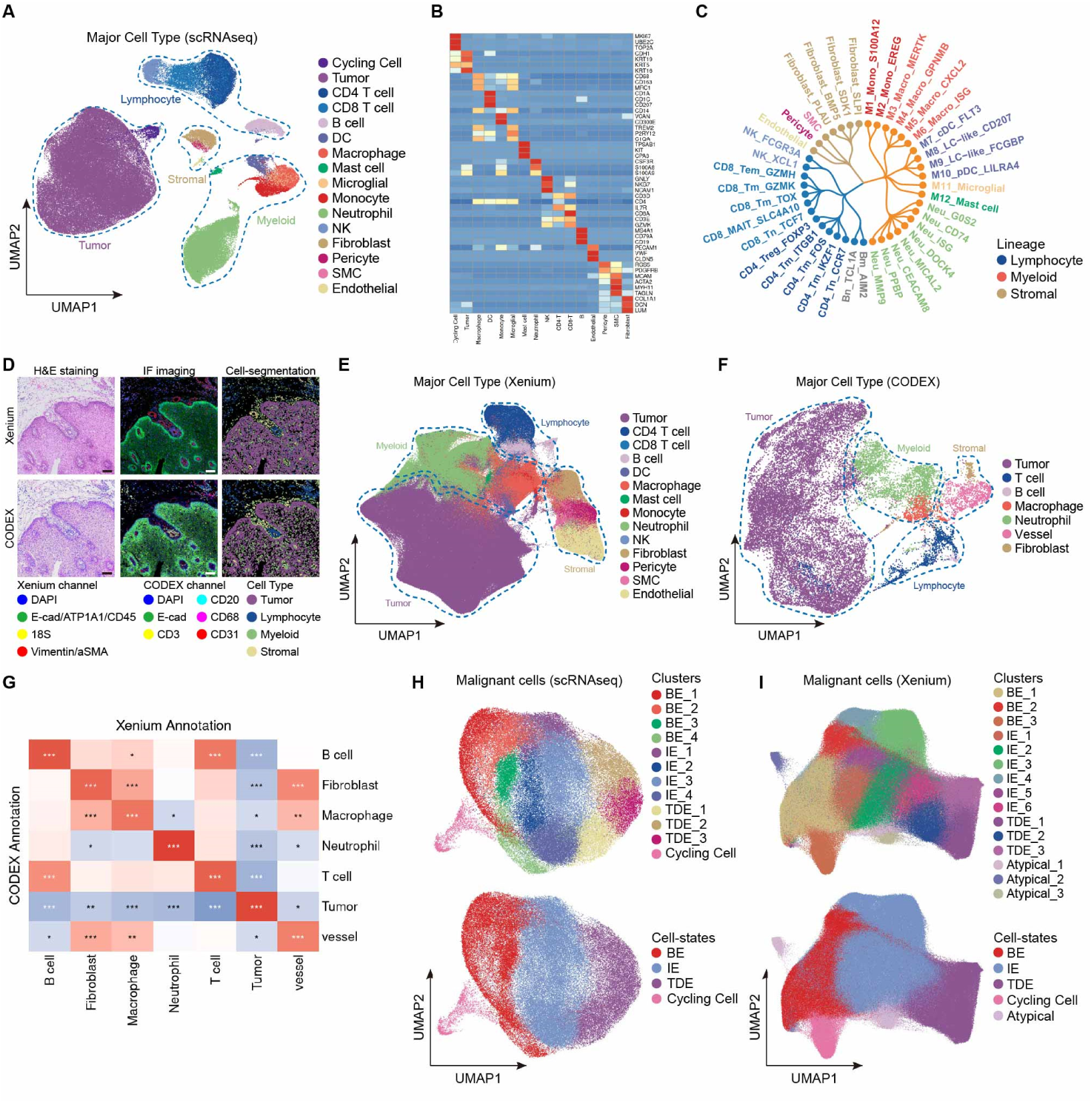
Multi-omics analysis identifies cell types in PCP. **A**: UMAP plot shows clusters in PCP tissues based on analysis of 10X GENOMICS scRNA-seq data. **B**: Heatmap showing the expression levels of marker genes in each scRNA-seq cluster. **C**: The branching tree diagram illustrates the annotations of lymphocyte, myeloid, and stromal cell subsets in PCP. **D**: Xenium and CODEX assays were performed on serial sections. H&E staining revealed the histological morphology of the same regions. Cell segmentation was performed based on immunofluorescence staining, and cell annotations were projected onto the digital pathological images. **E**: UMAP plot shows clusters in PCP tissues based on analysis of Xenium data. **F**: UMAP plot shows clusters in PCP tissues based on analysis of CODEX data. **G**: Heatmap shows the correlation of cell proportions for each cell type in Xenium and CODEX data. **H**: UMAP plot shows tumor subclusters and cell-states based on analysis of scRNA-seq data. **I**: UMAP plot shows tumor subclusters and cell-states based on analysis of Xenium data.

To further delineate the spatial molecular landscape of PCP, we performed spatial transcriptomic and spatial proteomic profiling on serial tissue sections (Fig. 1D). First, using the 10x Genomics Xenium platform, we profiled a 5,000-gene panel at single-cell resolution. As a result, we obtained the transcriptomes of 1,738,658 cells and performed clustering and cell type annotation (Fig. 1E). In parallel, we applied CODEX multiplexed fluorescence imaging to serial sections, enabling the spatial quantification of 27 proteins across 1,640,750 cells (Fig. 1F). The relative abundances and absolute numbers of major cell types showed significant concordance between patient-matched Xenium and CODEX datasets (Fig. 1G).

We next performed a more granular analysis of the tumor cell compartment to define the molecular phenotypes and functional heterogeneity of PCP tumor cells. Based on single-cell transcriptional profiles, we identified four recurrent cell-states of tumor cells that coexisted across patients: basal-like epithelial state (BE), intermediate epithelial state (IE), terminally differentiated epithelial state (TDE), and cycling cell (CC) (Fig. 1H). These cell-states were characterized by distinct molecular markers and functional programs, revealing substantial intratumoral heterogeneity within PCP (Fig. S1). Consistently, reclustering of tumor cells in the Xenium dataset identified cell-states and marker genes that closely recapitulated those observed in the scRNA-seq data (Fig. S1). Moreover, the antibody panel designed for CODEX imaging enabled the identification of the major tumor cell-states using a limited combination of protein markers. Collectively, our spatially resolved multi-omics cellular atlas delineated four transcriptionally distinct cell-states of tumor within PCP, providing a refined framework for understanding tumor cell heterogeneity in this disease.

## Discussion

Papillary craniopharyngioma (PCP) is a histologically low-grade central nervous system tumor, yet its clinical management remains particularly challenging. PCPs typically arise in the sellar and suprasellar regions and are closely associated with critical neuroanatomical structures, including the hypothalamus, pituitary stalk, optic chiasm, and third ventricle. Surgical resection therefore requires a delicate balance between achieving durable tumor control and preserving neurological and neuroendocrine function. Although more extensive resection may improve local tumor control, it can also cause irreversible injury to the hypothalamic–pituitary axis, resulting in long-term complications such as hypopituitarism, diabetes insipidus, severe obesity, sleep–wake disturbances, cognitive impairment, and hypothalamic syndrome. These persistent neuroendocrine and metabolic sequelae substantially compromise postoperative quality of life, underscoring the limitations of surgery as the sole strategy for long-term management of PCP.

In recent years, the identification of the BRAF V600E mutation and the introduction of BRAF/MEK-targeted therapy have represented major advances in the precision treatment of PCP. Nevertheless, compared with malignancies such as melanoma and lung cancer, for which comprehensive molecular classifications and tumor microenvironmental frameworks have been established, the fundamental biology of PCP remains poorly characterized. Previous studies have largely focused on its histopathological features and relatively simple core genetic driver, whereas the diversity of tumor cell states, functional differences among tumor populations, and their dependence on the surrounding microenvironment remain insufficiently understood. This knowledge gap limits our ability to explain the heterogeneity in PCP growth patterns, clinical behavior, and therapeutic responses, and has hindered the development of precision interventions directed against specific tumor cell states or microenvironment-dependent mechanisms. Thus, even in the presence of a well-defined molecular target, a deeper understanding of intratumoral heterogeneity and the tumor microenvironment may be essential for advancing PCP treatment from a predominantly driver mutation–oriented paradigm toward cell-state- and niche-informed precision therapy.

In this study, we systematically characterized PCP heterogeneity across multiple complementary dimensions by integrating single-cell transcriptomics, single-cell spatial transcriptomics, and high-dimensional spatial proteomics. At the single-cell transcriptomic level, we found that PCP is not composed of a transcriptionally homogeneous tumor cell population, but instead comprises multiple stable and recurrent tumor cell states. Specifically, we identified four major tumor cell states: BE, IE and TDE. Each cell-state was characterized by distinct molecular markers and transcriptional programs and was reproducibly observed across multiple patients, suggesting that these states are unlikely to reflect patient-specific variation or stochastic transcriptional fluctuations, but rather represent recurrent and biologically relevant functional states within PCP.

These findings extend the conventional histopathological view of PCP tissue architecture. PCP has traditionally been regarded as a tumor with relatively uniform epithelial morphology and a comparatively simple genetic background. Our data, however, indicate that genetic simplicity does not equate to cellular homogeneity. Even in the context of a shared core oncogenic driver, PCP tumor cells can occupy distinct differentiation, proliferative, and functional states, thereby generating substantial phenotypic heterogeneity. Compared with approaches that characterize tumors solely on the basis of mutational status or bulk tissue expression profiles, a cell-state-based framework provides a more refined view of the functional diversity of tumor cell populations within PCP.

The distinct molecular features of BE, IE, and TDE further suggest the existence of a continuous differentiation hierarchy among PCP tumor cells. BE cells display prominent basal-like and less differentiated features, IE cells exhibit molecular programs intermediate between basal-like and terminally differentiated states, whereas TDE cells show a more mature and terminally differentiated epithelial phenotype. These populations may therefore represent dynamic cellular states along a differentiation continuum rather than completely independent cell types. Consistent with this interpretation, complementary analyses using CytoTRACE 2 and RNA velocity can provide orthogonal information on developmental potential and transcriptional dynamics, respectively, thereby supporting potential state-transition relationships among these tumor cell populations. Although such transcriptome-based inference cannot substitute for direct lineage tracing, it provides a conceptual framework for understanding how PCP tumor cells may transition across distinct differentiation states.

By integrating single-cell transcriptomics, single-cell spatial transcriptomics, and spatial proteomics, our study reveals the multilayered heterogeneity of PCP across tumor cell states, microenvironmental composition, and spatial organization. Despite its relatively restricted set of core genetic driver events, the tissue ecosystem of PCP is substantially more complex than previously appreciated. Distinct tumor cell states coexist with diverse immune and stromal populations to form a highly organized spatial tumor ecosystem. These findings extend the biological understanding of PCP beyond conventional histopathology and driver mutations toward a framework centered on tumor cell states and the spatial microenvironment. This framework provides a biological basis for investigating disease progression, interpatient variability in therapeutic response, and potential mechanisms of treatment resistance, while also laying the groundwork for the development of precision therapeutic strategies targeting specific tumor cell states and microenvironmental niches.

## Materials and Methods

### 1. Human tissue sample collection

This study was approved by the Ethics Committee of West China Hospital, Sichuan University, and the Ethics Committee of Sun Yat-sen University Cancer Center (approval no. 20211047A). All PCP samples included in the discovery cohort were collected at West China Hospital, Sichuan University, and written informed consent was obtained from all participants.

A total of 45 patients with papillary craniopharyngioma (PCP) were enrolled in the discovery cohort. In addition, 29 patients with PCP treated at Sun Yat-sen University Cancer Center were included as an independent validation cohort. All study procedures were conducted in accordance with the principles of the Declaration of Helsinki.

### 2. Tissue collection and processing

Freshly resected PCP tumor specimens were processed immediately after surgical removal according to the requirements of the corresponding downstream multi-omics assays.

For single-cell RNA sequencing (scRNA-seq), tumor tissues were mechanically minced and enzymatically dissociated into single-cell suspensions. The resulting suspensions were filtered to remove tissue debris and cell aggregates, and low-quality or nonviable cells were excluded before library preparation. Peripheral blood mononuclear cells (PBMCs) were isolated from peripheral blood samples by Ficoll density-gradient centrifugation.

For Xenium spatial transcriptomic and CODEX spatial proteomic analyses, tumor specimens were processed according to the respective platform-specific workflows, including fixation, embedding, sectioning, and tissue pretreatment. Serial tissue sections were used for Xenium and CODEX profiling to minimize the impact of intratumoral spatial heterogeneity on cross-platform comparisons and to facilitate spatially matched multi-omics analyses.

### 3. Single-cell RNA sequencing

Single-cell suspensions derived from PCP tumor tissues and matched peripheral blood mononuclear cells (PBMCs) were used for library preparation and sequencing using the 10x Genomics single-cell RNA sequencing platform. Fresh PCP tumor tissues were dissociated into single-cell suspensions according to the manufacturer’s recommended protocol, and only suspensions with cell viability greater than 90% were used for subsequent library construction. PBMCs were isolated from peripheral blood collected from patients with PCP by Ficoll density-gradient centrifugation and processed to obtain high-viability single-cell suspensions.

Single-cell suspensions were loaded onto the Chromium Single Cell Controller (10x Genomics) to generate single-cell gel beads in emulsion (GEMs). Approximately 20,000 cells were loaded into each channel, with a target recovery of 5,000–10,000 cells per sample. Reverse transcription, post-reverse-transcription cleanup, cDNA amplification, and library construction were subsequently performed according to the manufacturer’s instructions. Library quality and concentration were assessed before sequencing. The resulting scRNA-seq libraries were sequenced on an Illumina NovaSeq 6000 platform to generate 150-bp paired-end reads.

### 4. Data integration, dimensionality reduction, clustering, and cell-type annotation

Downstream analysis of scRNA-seq data was performed using the Seurat package. Gene expression matrices were normalized using either the NormalizeData or SCTransform function, as appropriate. To integrate data from multiple patients and minimize sample-specific batch effects, the IntegrateLayers function in Seurat was applied using Harmony-based integration.

For dimensionality reduction and unsupervised clustering, the integrated expression matrix was scaled using ScaleData, followed by principal component analysis (PCA) using RunPCA. The first 50 principal components were initially calculated, and the first 25 principal components were subsequently used to construct the shared nearest-neighbor graph and perform unsupervised clustering using FindNeighbors and FindClusters, respectively. The resulting cell clusters were visualized in two-dimensional space using Uniform Manifold Approximation and Projection (UMAP) or t-distributed stochastic neighbor embedding (t-SNE).

Cluster-specific marker genes were identified using the FindAllMarkers function implemented in Seurat. Cell-type identities were assigned by integrating annotation results generated using the SingleR package with the expression patterns of canonical lineage-specific marker genes reported in previous studies. Cell-type annotations were further manually reviewed on the basis of established marker expression to ensure biological consistency.

### 5. Xenium In Situ spatial transcriptomic profiling

Xenium In Situ RNA imaging was performed using the 10x Genomics Xenium platform. To minimize sampling bias arising from intratumoral spatial heterogeneity, tissue microarrays (TMAs) were constructed from PCP tumor specimens. Multiple representative regions were sampled from each tumor, with each tissue core measuring 2 mm in diameter. The final TMA comprised 72 spatially independent tissue spots from 24 patients, with an average of three spots per patient.

Tissue sections approximately 10 μm in thickness were prepared and mounted within the designated imaging area of Xenium slides. After drying, the sections were fixed with paraformaldehyde (PFA) and subsequently permeabilized with 70% ethanol according to the manufacturer’s recommended workflow. Spatial transcript detection was performed using the Xenium 5K gene panel. Following tissue pretreatment, target-specific RNA probes were hybridized to their corresponding transcripts, followed by probe ligation and signal amplification according to the standard Xenium In Situ workflow. Tissue autofluorescence was subsequently quenched, and nuclei were stained to facilitate tissue visualization, single-cell segmentation, and downstream assignment of detected transcripts to individual cells. Prepared slides were loaded onto the Xenium Analyzer for automated iterative imaging and in situ transcript decoding. Multiple rounds of fluorescence imaging were used to identify individual RNA molecules, with both gene identity and two-dimensional spatial coordinates recorded for each detected transcript. Nuclear and tissue morphology images were acquired simultaneously to support subsequent cell segmentation. Following whole-slide imaging, the TMA regions were manually inspected, and regions of interest (ROIs) were selected based on tissue integrity, tumor representation, and imaging quality. Each 2-mm TMA spot was treated as an independent spatial region for downstream analyses, while patient identity and the correspondence among multiple spots derived from the same patient were retained to account for both interpatient and intrapatient spatial heterogeneity. Post-run data processing was performed using the default Xenium analysis pipeline, including image processing, transcript decoding, nuclear identification, cell segmentation, and assignment of transcripts to individual cells. The resulting dataset contained single-cell gene expression profiles, spatial coordinates, cell segmentation information, and transcript-level spatial localization, and was subsequently used for cell-type annotation, tumor cell-state identification, and spatial microenvironment analyses.

### 6. CODEX multiplexed imaging

To characterize the spatial proteomic landscape of PCP at single-cell resolution, multiplexed imaging was performed using the CODEX platform. A curated panel of oligonucleotide-conjugated antibodies targeting tumor, immune, and stromal cell markers was first validated for specificity and staining performance. Following panel validation, a pilot CODEX run was performed to optimize the signal-to-noise ratio, antibody dilution, fluorescence exposure time, and imaging cycle assignment for each antibody conjugate. Multiplexed CODEX imaging was subsequently performed using a cyclic workflow consisting of sequential stripping, annealing, and fluorescence imaging of reporter oligonucleotides complementary to the unique oligonucleotide barcodes conjugated to individual antibodies. Multiple imaging cycles were conducted to enable spatial detection of the complete antibody panel within the same tissue section while preserving tissue architecture and single-cell spatial information.

Following completion of multiplexed imaging, raw QPTIFF images were imported into QuPath (version 0.3.2) for initial image processing and quality control. Image preprocessing included image stitching, correction for inter-cycle spatial drift, deconvolution, and concatenation of individual imaging cycles. Regions affected by imaging artifacts or compromised tissue morphology, including out-of-focus areas, tissue folds, and debris, were manually annotated and excluded from subsequent analyses. Single-cell segmentation was performed using the deep-learning-based StarDist algorithm applied to the DAPI nuclear channel with default model parameters. Prior to segmentation, images were downsampled by 50%, and contrast-limited adaptive histogram equalization was applied to enhance nuclear contrast and improve segmentation performance. Following nuclear segmentation, individual cell boundaries were defined and quantitative protein expression measurements were extracted for each segmented cell. The spatial centroid of each cell was determined from its x-y coordinates, thereby generating a single-cell dataset containing both multiplexed protein expression profiles and spatial localization information. The quality of cell segmentation was further evaluated by experienced pathologists through visual inspection of representative tissue regions to ensure accurate identification of individual cells and adequate correspondence between segmentation boundaries and tissue morphology. Cells and tissue regions that did not meet predefined image-quality or segmentation criteria were excluded from downstream analyses.

### 7. RNA velocity analysis

RNA velocity analysis was performed to infer the directionality and dynamics of transcriptional state transitions among PCP tumor cells. Spliced and unspliced transcript abundances were quantified from the 10x Genomics scRNA-seq alignment data using velocyto and subsequently analyzed using the scVelo framework. A velocity graph was constructed on the basis of the consistency between estimated RNA velocity vectors and the transcriptional relationships among neighboring cells. Velocity streamlines were subsequently projected onto the UMAP representation to visualize directional transitions among BE, IE, TDE, and other tumor cell states. In addition, latent time was inferred using the dynamical model to provide an internally reconstructed temporal ordering of tumor cells based on their transcriptional kinetics. The inferred velocity trajectories and latent-time distributions were jointly evaluated with CytoTRACE 2 potency scores to characterize the differentiation hierarchy and potential state-transition relationships among PCP tumor cell populations.

### 8. Statistical analysis

Data statistics are performed based on GraphPad Prism 8.0 software. We obtained the p value of the difference between different groups by performing Student’s t-test and one-way ANOVA. When p < 0.05, differences were considered statistically significant.

## 9. Data availability

The scRNA-seq, CODEX and Xenium data reported in this paper have been deposited in the OMIX, China National Center for Bioinformation/Beijing Institute of Genomics, Chinese Academy of Sciences (https://ngdc.cncb.ac.cn/omix, accession no. OMIX021470, OMIX021471 and OMIX021475). All relevant data supporting this study are available from the corresponding author upon reasonable request.

